# High-Contrast PET imaging with [^18^F]-NT160, a Class-IIa Histone Deacetylase (Class-IIa HDAC) Probe for In Vivo Imaging of Epigenetic Machinery in the Central Nervous System

**DOI:** 10.1101/2022.11.12.516260

**Authors:** Nashaat Turkman, Sulan Xu, Chun-Han Huang, Christopher Eyermann, Julia Salino, Palwasha Khan

## Abstract

We utilized positron emission tomography (PET) imaging *in vivo* to map the spatiotemporal biodistribution/expression (protein density) of class-IIa histone deacetylases (class-IIa HDACs) in the brain. Herein, we report an improved radiosynthesis of [^18^F]-NT160 using 4-hydroxy-TEMPO which led to a significant improvement in radiochemical yield and molar activity. PET imaging with [^18^F]-NT160, a highly potent class-IIa HDAC inhibitor with sub-nM affinity for HDAC4 and 5 isoforms, led to high-quality and high-contrast images among various brain regions. [^18^F]-NT160 displayed excellent pharmacokinetic and imaging characteristics: brain uptake is high in gray matter regions, leading to high-quality PET images; tissue kinetics are appropriate for an ^18^F tracer and specific binding for class-IIa HDACs is demonstrated by self-blockade. Higher uptake with [^18^F]-NT160 was observed in the hippocampus, thalamus, and cortex while there was relatively lower uptake in the cerebellum and striatum. Overall, our current studies with [^18^F]-NT160 will likely facilitate the development and clinical translation of class-IIa HDACs of the next generation of PET tracers for imaging and targeted therapy of cancer and the diseases of the central nervous system (CNS).

## Introduction

The histone deacetylases (HDACs) are a family of 18 enzymes and the class-IIa HDACs is a sub-family of four members (HDACs: 4, 5, 7 and 9). The dysregulation of class-IIa HDAC in the brain has been implicated in human cancers ^1–3^ and the disorders of the central nervous system (CNS) such as stroke ^4–7^, Huntington’s ^8–10^ and Alzheimer’s diseases ^11–13^. However, non-invasive imaging biomarkers for quantifying the class-IIa HDAC expression in tumors and in the CNS are lacking. Therefore, we are targeting class-IIa HDACs as a novel theranostic for molecular imaging and targeted therapy. The aim of our ongoing work is to develop positron emission tomography (PET) radiotracers specific to class-IIa HDAC for non-invasive quantification and mapping of the biodistribution of class-IIa HDAC protein expression in cancer and brain diseases.

Initial efforts to develop radiolabeled HDAC inhibitors were hampered by poor blood-brain barrier (BBB) permeability^14–17^ which limited their utility for brain imaging. [^11^C]-martinostat was the first successful brain penetrant class-I HDAC (HDAC1, 2 and 3) inhibitor-based tracer^18–20^. Also, PET imaging in rodent, nonhuman primates, and human subjects was reported with a brain-penetrant and semi-selective inhibitor of HDAC6 (class-IIb HDAC) ^21, 22^. Furthermore, class-IIa HDACs substrate-based radiotracers exhibited fast unspecific degradation and poor pharmacokinetic *in vivo* that were detrimental to their clinical utility as PET radiotracers ^23^. The main limitation to the utility of the substrate-based radiotracers is that the radiolabeled metabolites were taken up by glial cells ^24, 25^ and the metabolite based radiotracers did not remain localized to the point of catabolism by class-IIa HDACs ^20^ thus, confounding the PET signal leading to inaccurate map of biodistribution of class-IIa HDAC in the brain. Therefore, the substrate-based radiotracers do not inform on neither expression nor the activity of class-IIa HDACs thus precluding their practical utility for quantitative imaging. To overcome these limitations, we developed an inhibitor-type radiotracers thus enabling accurate quantification of class-IIa HDACs protein expression/density in the brain by PET imaging.

Our publication provides the first accurate and quantitative information on the biodistribution/expression (protein density) of class-IIa HDACs in the CNS. Class-IIa HDACs are known to be highly expressed in various regions of the brain and their overexpression has been indicated in various brain disorders ^26, 27^. Therefore, mapping the biodistribution of class-IIa HDACs in the brain will pave the way for quantitative measurement of their expression in the healthy brain compared to the disease state.

## Results and Discussion

We recently reported, the design synthesis and structure activity relationship (SAR) campaign of trifluoromethyl-oxadiazole (TFMO) containing molecules and identified a highly selective and potent small molecule inhibitors of class-IIa HDACs suitable for PET tracer development ^28^. We also reported a late stage radiosynthesis of the TFMO moiety with [^18^F]-fluoride ^29^ and we utilized the methodology to radiosynthesize [^18^F]-TMP195 and [^18^F]-NT160 (Figure 1) and demonstrated their brain entry in mice ^28^. However, the low spatial resolution of the uPET does not permit obtaining quantitative spatiotemporal biodistribution of class-IIa HDAC in the brain. Therefore, to overcome this limitation, we performed PET imaging studies, biodistribution, and metabolite studies with [^18^F]-NT-160 in Sprague Dawley (SD) rats.

**Figure 1.**
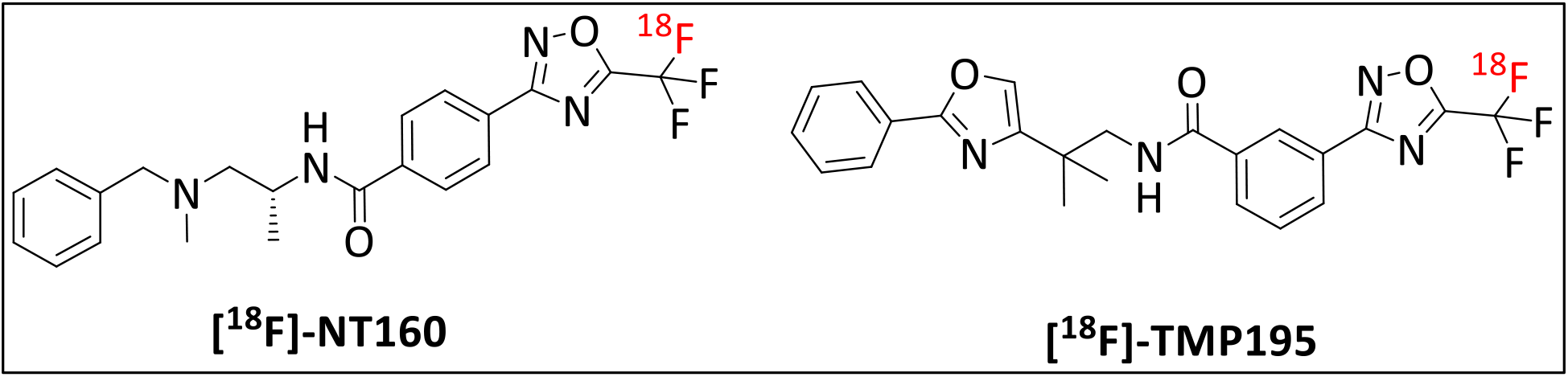
Chemical structures of radiolabeled class-IIa HDAC inhibitors.

### Improved radiosynthesis of [^18^F]-NT160 with the aid of 4-hydroxy-TEMPO

The radiosynthesis of [^18^F]-NT160 was performed similar to our previous report ^28, 29^ with modifications. Performing the radiochemical reaction at 175 ^o^C with the addition of a small quantity (0.3-0.5 mg) of 4-hydroxy-2,2,6,6,-tetramethylpiperidine-1-oxyl (known as 4-hydroxy-TEMPO or TEMPOL) led to a significant improvement of the radiochemical yield of 7.5 ±1.0%, (decay corrected) and the molar activity of 0.55–0.96 GBq/umol (15–26 mCi/umol) compared to the previous work: radiochemical yield of 2-5% (decay corrected) and the molar activity of 0.33–0.49 GBq/umol (8.9–13.4 mCi/umol) ^28^.

**Scheme 1.**
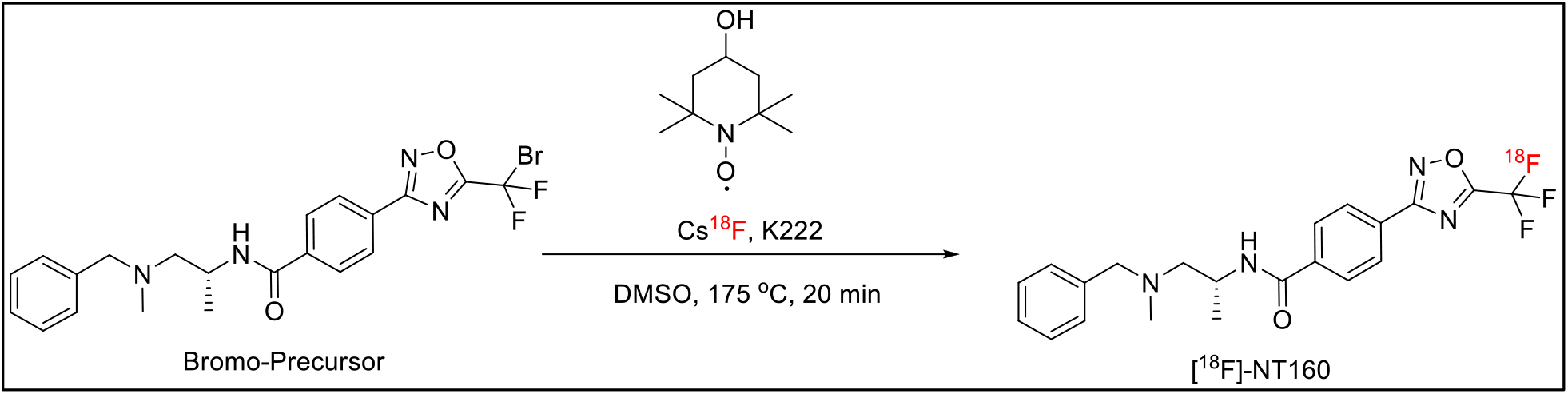
Improved radiosynthesis of [^18^F]-NT160

The use of 4-hydroxy-TEMPO allowed for performing the reaction at 165 °C, a relatively higher temperature than 150 ^o^C used in our previous report^28, 29^. Interestingly, increasing the radiochemical reaction to 175-200 ^o^C produced similar radiochemical yields but with significantly reduced molar activity. In contrast, performing the radiochemical reaction at <160 ^o^C led to a significantly reduced radiochemical yield. We initially hypothesized that 4-hydroxy-TEMPO protected the bromo-precursor by quenching the free radicals present or released into the reaction mixture at high reaction temperature. However, remarkably, the use of TEMPO, which is also a free radical quencher, did not produce improvement in the radiochemical yield or the molar activity which suggests that other mechanism might be involved in improving the radiochemical yield and the molar activity. The bromo-precursor remained mostly intact during the radiosynthesis despite the high temperature which suggests that 4-hydroxy-TEMPO is key to protect reactants from decomposition at high temperatures. Overall, this improvement is highly significant since it cut the amount of nonradioactive mass dose of the NT160 by >50% thus enabling performing blocking (target engagement) studies *in vivo* and will likely enable clinical translation of [^18^F]-NT160 by starting with larger amounts of radioactivity (i.e. >3.0 Ci). We also recommend the general utility of 4-hydroxy-TEMPO for improving the yields of radiochemical reactions that require high temperatures with specific emphasis on the ^18^F-trifluoromethyl moiety.

### Formulation and *in situ* stability of [^18^F]-NT160

[^18^F]-NT160 was formulated in 20% ethanol, 20% polysorbate 80 and 60% saline containing sodium ascorbate (5mg/mL). [^18^F]-NT160 remained intact (stable) with >98% radiochemical purity up to four hours post formulation based on the analysis with high performance liquid chromatography (HPLC).

### PET/CT Imaging

PET and PET/CT fusion with [^18^F]-NT160 produced high-quality and high-contrast images among various brain regions. [^18^F]-NT160 displayed excellent pharmacokinetic and imaging characteristics: brain uptake is high in gray matter regions, leading to high-quality PET images; tissue kinetics are appropriate for an ^18^F tracer and specific binding for class-IIa HDAC is demonstrated by self-blockade. Higher uptake with [^18^F]-NT160 was observed in the hippocampus, thalamus, and cortex, with lower uptake in the striatum and cerebellum.

High contrast is demonstrated between the brain and the surrounding tissues (Figure 2A-F). Our PET data indicates that [^18^F]-NT160 exhibited excellent *in vivo* characteristics, with rapid BBB penetration and observable washout pharmacokinetics. The brain uptake in the brain peaked at ~1.25 %ID/cc (SUV = 3.2) in the first five min thus exceeding the benchmark (>0.1%ID/cc) for successful CNS PET tracers ^30^. Remarkably, no indication of defluorination (no radioactive uptake in skull bone) was observed (Figure 2B and D). Defluorination leads to the fluoride ion accumulating in the bone of the skull, which can lead to the signal “spilling” into the brain thus confounding the quantification of changes in signal in cortical regions ^31, 32^.

**Figure 2.**
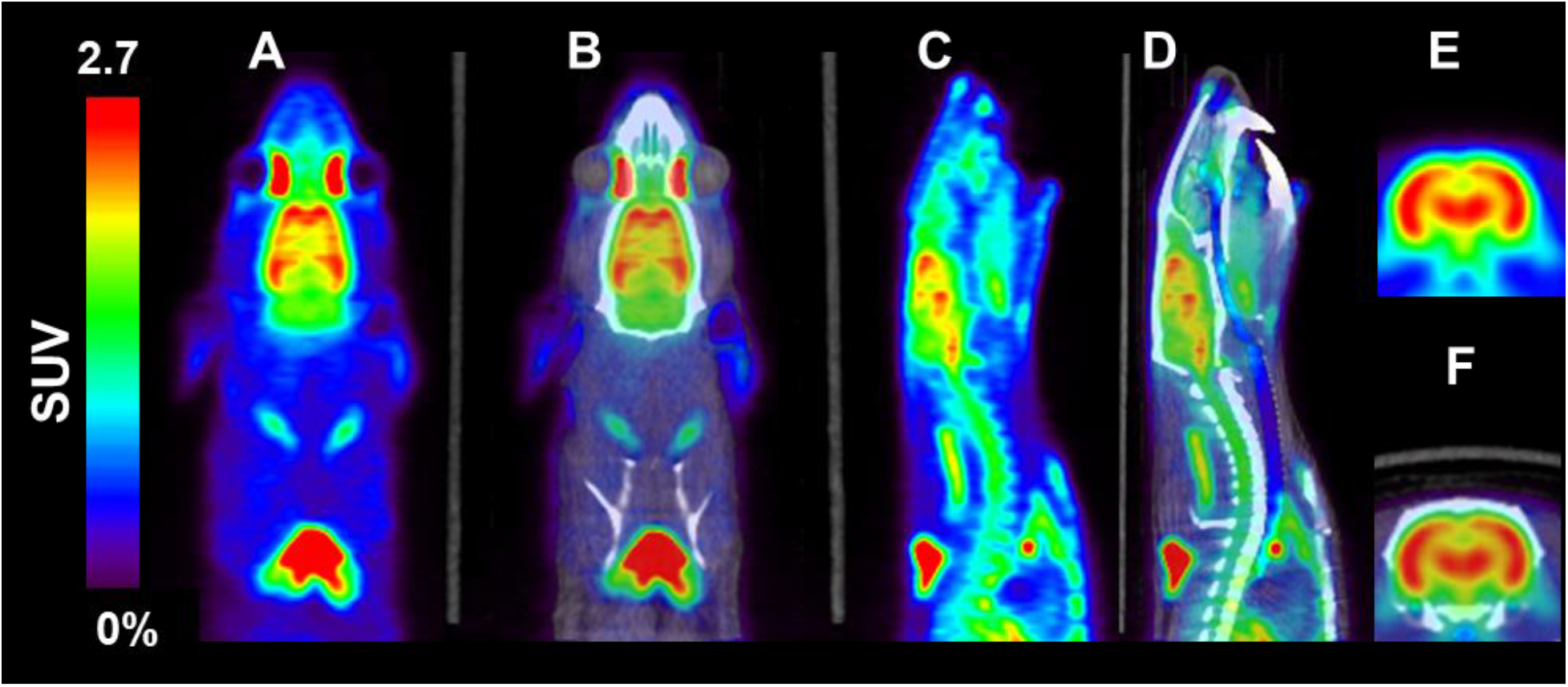
Representative PET and PET/CT (fusion) images with [^18^F]-NT160 of rat brain (summed 5-30 min): A) and B) coronal; C) and D) sagittal and E) and F) transverse respectively.

High brain-to-heart and brain-to-muscle ratios were observed with [^18^F]-NT160 at ten minutes post injection and were maintained for the rest of the scan time (Figure 3A and Table 2). Time–activity curves generated from the brain and heart signals showed that [^18^F]-NT160 cleared rapidly from the heart and the tracer uptake stabilized at 10 min post injection with an SUV = 0.4 (Figure 3A). Additionally, [^18^F]-NT160 exhibited relatively low uptake in the muscle, which peaked and plateaued at an SUV = 0.6. The uptake of [^18^F]-NT160 in muscle is likely specific and was displaceable (Figure 4: S.I.), which is consistent with the previous reports that class-IIa HDACs were expressed in the skeletal muscle ^33, 34^. However, the uptake in the periphery is likely confounded by the non-specific uptake from the radiometaolite fraction in the blood which constitutes the major radioactive fraction in the blood (discussed in the radiometabolism section below). Therefore, it is unclear as to whether the radiometabolite in the blood also contributed to the uptake in the muscle. Nevertheless, our imaging data demonstrate for the first time that the protein density/expression of class-IIa HDACs in the normal/healthy muscle is low when compared to the brain.

**Figure 3.**
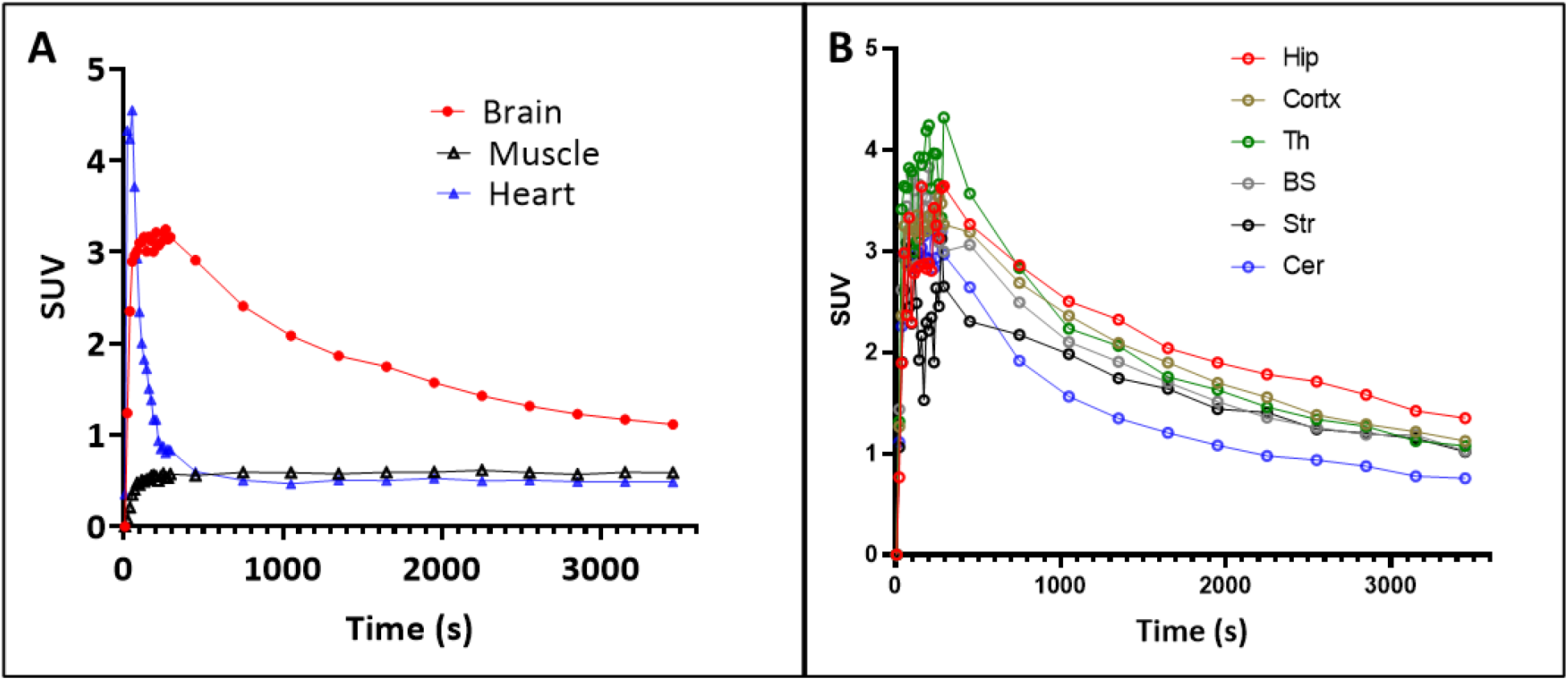
Time–activity curves of [^18^F]-NT160 obtained from dynamic imaging for 60 min. A) Brain, muscle, and heart; B) hippocampus (Hip), Cortex (cortx), Thalamus (TH), Striatum (Str) and cerebellum (Cer).

**Figure 4.**
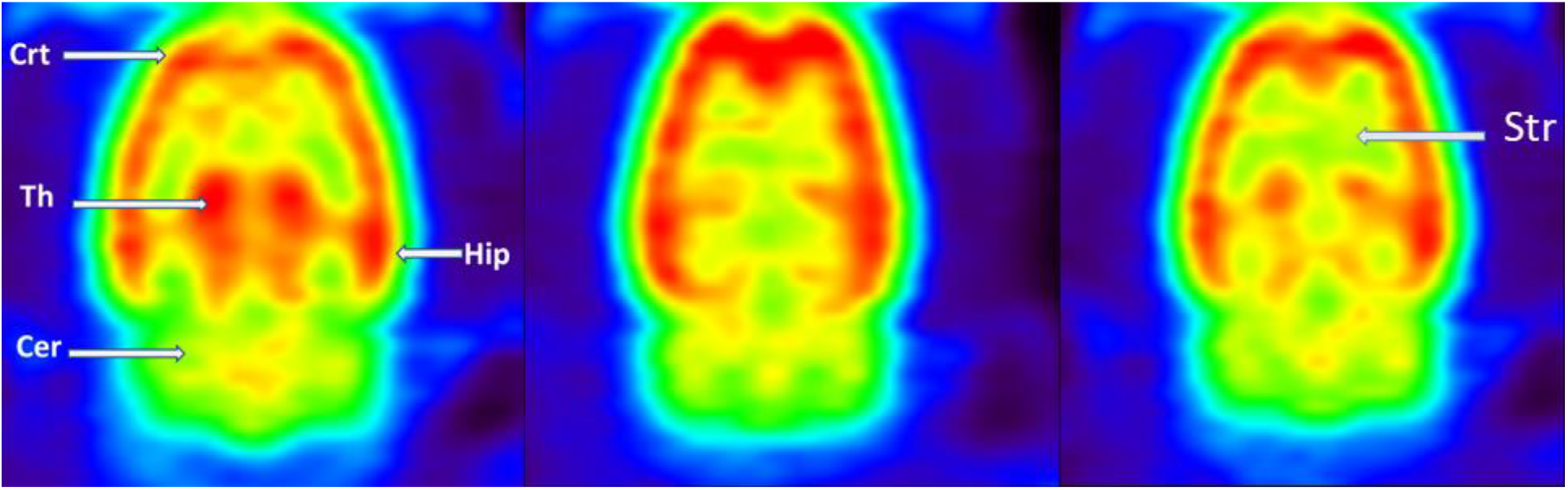
Representative coronal PET images (summed, 5-30 min) with [^18^F]-NT160 with the corresponding rat brain regions. Crt: Cortex, Cer: cerebellum, Th: thalamus, Hip: hippocampus, Str: striatum. The alignment for the images with the rat brain atlas was used to identify these regions of interest ^35, 36^.

**Table 1.** Radiochemical reaction conditions, radiochemical yields (RCY) and molar activity

| Temperature | 4-hydroxy-TEMPO (mg) | %RCY | Molar activity (A<sub>m</sub>) |
| --- | --- | --- | --- |
| 175 °C | 0.3-0.5 | 7.5 ±1.0 | 0.74–1.51 GBq/umol (20–41 mCi/umol) |
| 175 °C | 0 | <1.0 | ND |
| 185-200 °C | 0.3-0.5 | 7.5 ±1.0 | 0.22–0.32 GBq/umol (6.0–8.4 mCi/umol) |
| 150 °C (previous work)<sup>28,29</sup> | 0 | 3.5 ±1.5 | 0.33–0.49 GBq/umol (8.9–13.4 mCi/umol) |

**Table 2.** Biodistribution of [^18^F]-NT160 (SUV ±SEM) in rat brain, heart, and muscle.

|  | Time after tracer injection |  |  |  |
| --- | --- | --- | --- | --- |
|  | 1 min | 10 min | 30 min | 60 min |
| **Brain** | 2.96 ±0.95 | 2.40 ±0.44 | 1.57 ±0.29 | 1.11 ±0.26 |
| **Heart** | 3.71 ±2.29 | 0.51 ±0.27 | 0.52 ±0.16 | 0.48 ±0.18 |
| **Muscle** | 0.40 ±0.29 | 0.59 ±0.17 | 0.59 ±0.15 | 0.59 ±0.15 |
| **Hippocampus** | 2.37 ±0.74 | 2.86 ±0.28 | 1.90 ±0.27 | 1.34 ±0.23 |
| **Cerebellum** | 3.01 ±1.08 | 1.91 ±0.89 | 1.08 ±0.25 | 0.76 ±0.18 |
| **Hippocampus-to-Cerebellum ratio** | 0.79 ±0.91 | 1.50 ±0.59 | 1.76 ±0.26 | 1.76 ±0.21 |
| **Brain-to-heart ratio** | 0.80 ±1.62 | 4.71 ±0.35 | 3.01 ±0.23 | 2.29 ±0.22 |
| **Brain-to-muscle ratio** | 7.40 ±0.62 | 4.01 ±0.31 | 2.66 ±0.22 | 1.88 ±0.21 |

Furthermore, the time–activity curves (Figure 3B) generated from dynamic PET imaging of the brain regions of interest showed that [^18^F]-NT160 displayed heterogeneous uptake in the brain with higher tracer uptake in the hippocampus, thalamus, cortex, and brain stem compared to the cerebellum. The high contrast was observed among brain regions at ten minutes post injection and was sustained for 60 minutes post injection. Altogether, these data strongly indicate that [^18^F]-NT160 displayed an excellent pharmacokinetic profile *in vivo* with high brain-to-heart, brain-to-muscle and hippocampus-to-cerebellum ratios were maintained throughout the entire 60 min of the scan (Table 2). Based on the time activity curves (Figure 2B), the highest contrast among brain regions can be obtained by performing PET scan in the 10-60 min time frame.

### Spatiotemporal Biodistribution of ^18^F-NT160 as a Surrogate Biomarker of Class-IIa HDACs Expression (Protein Density) in the Brain

PET imaging (Figure 4) with [^18^F]-NT160 showed a high contrast among various regions of the brain with higher uptake was observed in the hippocampus, thalamus, brainstem and cortex and relatively lower uptake in the striatum and cerebellum. Our *in vivo* findings provide the first accurate and quantitative map of the biodistribution/expression (protein density) patterns of class-IIa HDACs in the CNS in its entirety.

Our data is consistent with previous reports on the heterogeneous biodistribution of class-IIa HDACs in the rodents’ brain ^37, 38^. However, the main limitation of the previous reports is the reliance on semi-quantitative or qualitative data to delineate the class-IIa HDAC biodistribution in the brain. HDAC4 was reported to be highly expressed in some mouse brain regions than in others and HDAC4 was absent from the white matter ^38^. Moreover, in situ hybridization which was used to determine the HDAC isoforms expression in >50 rat brain regions ^37^, showed that HDAC11, −3, −5 and −4 are expressed most highly, and HDAC10, −9, and −7 have the lowest expression levels throughout the brain. In agreement with our imaging data in rats, the expression of class-IIa HDAC4 and 5 were high in the rat cerebral cortex, thalamus, and hippocampus. In the cerebellum, HDAC4 and 5 expression was limited solely to the granule cell layer which can explain the relatively low tracer accumulation in the cerebellum. PET does not detect subcellular distribution of proteins but rather the protein density distribution in a specific region. Therefore, it is likely that the protein density class-IIa HDAC is less in the cerebellum compared to the hippocampus, thalamus, and brain stem. These results are in line with our *in vitro* autoradiography and histology with HDAC4 (Figure 5C). It is important to emphasize that PET is quantitative and more accurate than the available data on class-IIa HDAC expression in the brain which underscore the significance of our approach to non-invasively provide quantitative information on the level of class-IIa HDAC *in vivo* in real time.

**Figure 5.**
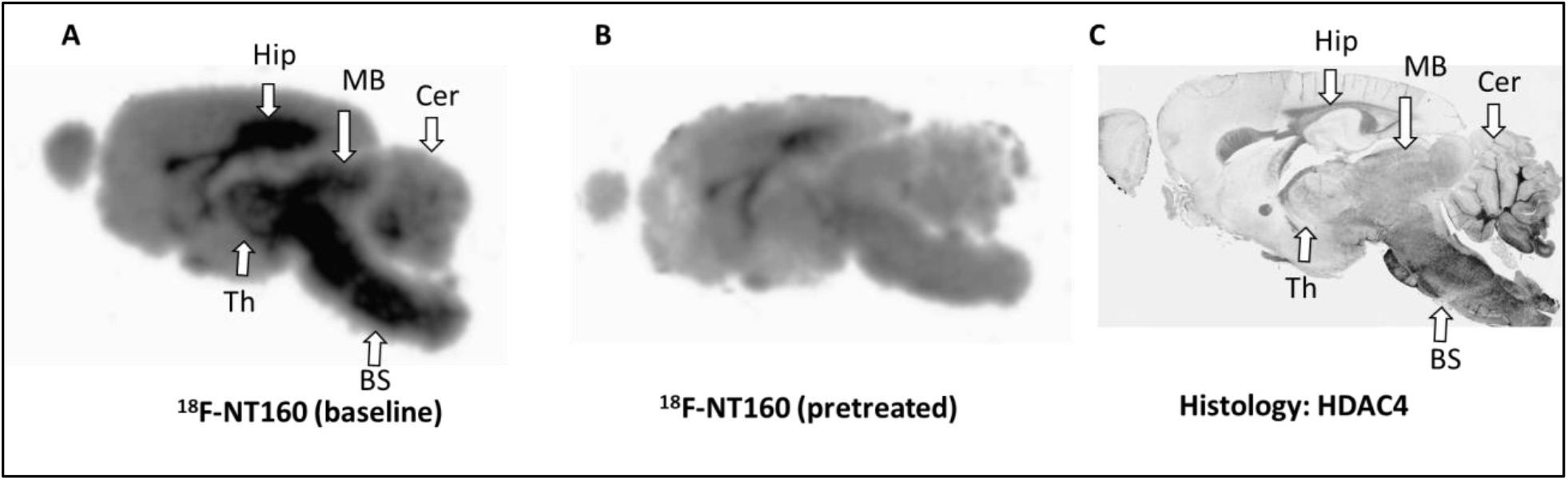
A) *In vitro* autoradiography with [^18^F]-NT160 at baseline and B) after pretreatment with NT160 (1.0 uM). C) Brain histology obtained with HDAC4 antibody.

### *In vitro* Autoradiography and Histology

We performed *in vitro* autoradiography with [^18^F]-NT160 and histology with HDAC4 (Figure 5). *In vitro* autoradiography with [^18^F]-NT160 showed that the highest uptake in the brain was observed in the hippocampus, mid brain, and brainstem with moderate uptake was observed in the rest of the brain. Brain uptake was significantly reduced by competitive blocking with NT160 (1.0 uM). Significant reduction in the tracer uptake was observed in the hippocampus, mid brain, and brain stem. Also, the tracer uptake was reduced to lesser extent in other brain regions. This data strongly indicates the brain uptake [^18^F]-NT160 was displaceable and specific to class-IIa HDACs. We also performed histology with HDAC4 antibody which also confirmed high HDAC4 expression in the hippocampus, mid brain, and brainstem (Figure 5). It is important to note that NT160 is somewhat selective for HDAC4 (~15 folds) over other class-IIa HDAC isoforms. Therefore, it is not surprising that the *in vitro* autoradiography with [^18^F]-NT160 is skewed toward HDAC4 and likely is representative of HDAC4 biodistribution in the brain. Preliminary histology with HDAC5 antibody showed high expression in the cortex and brain stem and lower expression was observed in the hippocampus.

### Specific Binding/Occupancy of [^18^F]-NT160 *In Vivo*

We initially performed the blocking experiment with [18F]-NT160 at 20 min post injection followed by side-by-side static PET imaging which was performed *ex-vivo* on the excised brains. Then, both brains were weighed, and the radioactivity was counted using a gamma counter. The brain uptake was reduced when comparing images obtained at the baseline to images obtained from the tracer co-administered with a dose of non-radioactive NT160 (1.0 mg/kg). Also, the brain uptake of [18F]-NT160, was reduced when counting the radioactivity (CPM: count per minute) using gamma counter and normalized to the brain weight (Figure 4, SI). Self-blocking with the unlabeled NT160 resulted in significant decrease in the total tracer uptake in the brain thus demonstrating specific binding of 18F-NT160 to class-IIa HDACs *in vivo.*

Moreover, brain uptake was also reduced *in vivo* (Figure 6) when comparing time-activity curves obtained from the tracer administered at the baseline with images obtained from the tracer co-admistered with 1.0 mg/kg of the non-radioactive NT160 (0.5 mg/kg in addition to 0.5 mg/kg calculated from the specific activity). Self-blocking with the unlabeled NT160 resulted in a significant decrease in the tracer uptake in the whole brain (30% reduction in the SUV) which demonstrates that that the binding/occupancy of [^18^F]-NT160 to class-IIa HDACs *in vivo* is specific and displaceable (Figure 6A). However, the overall decrease in radioactivity in the whole brain is rather simplistic, therefore, kinetic modeling of the PET data will be needed to accurately quantify the decrease in the nondisplaceable binding potential (BP_ND_) which is more informative of the extent of tracer binding/displacement. Also, as was discussed above, the expression/density of class-IIa HDAC in the brain is heterogeneous and therefore, regional displacement is more informative on the extent of class-IIa HDAC biodistribution in the brain than the displacement in the whole brain.

**Figure 6.**
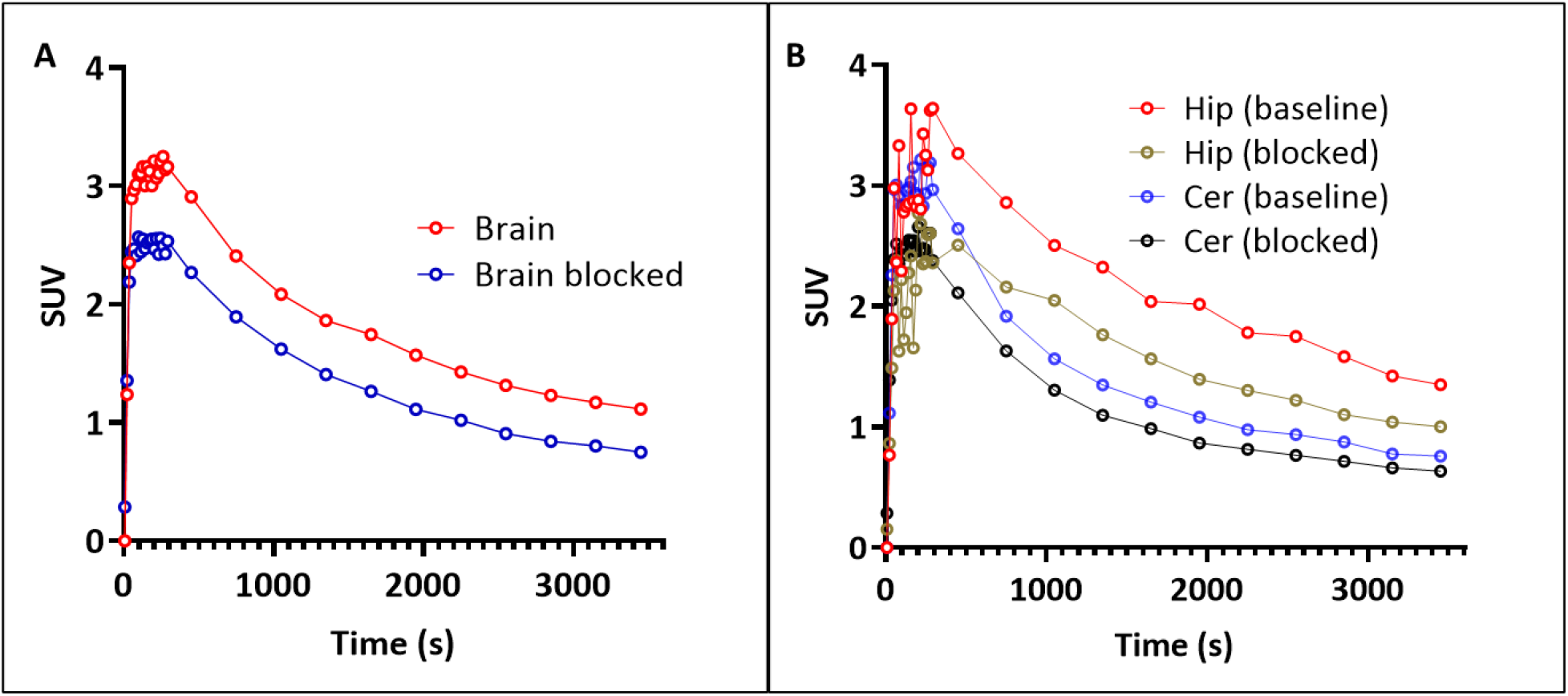
Time–activity curves obtained from dynamic imaging for 60 min: A) whole brain; B) hippocampus and cerebellum at baseline and after self-blocking, Cer: cerebellum, Hip: hippocampus.

The highest tracer uptake and consequently the largest reduction in tracer uptake due to blocking was observed in the hippocampus thus indicating high expression/density of class-IIa HDACs (Figure 6B). In contrast, the uptake of [^18^F]-NT160 was lowest in the cerebellum and the uptake was somewhat displaceable, however to a lesser ratio than the hippocampus (Figure 6B). Previous literature reports point to expression of class-IIa HDACs in the cerebellum localized to the Purkinje cells in the granule cell layer ^37, 38^. The higher reduction in tracer uptake in hippocampus by blocking is reflective of high occupancy and high density of class-IIa HDACs in the hippocampus. In contrast, the lower uptake and lower reduction in tracer uptake in the cerebellum is reflective of the low density of class-IIa HDACs in the cerebellum. Our *in vivo* findings are supported with *in vitro* autoradiography which showed high [^18^F]-NT160 accumulation in the hippocampus relative to the cerebellum (Figure 5A). Also, histology with HDAC4 antibody further supports the *in vivo* findings. Therefore, it is unclear whether the cerebellum can be utilized as a reference region for future kinetic modeling and quantification of the PET data. The tracer uptake in the cerebellum seems to be heterogeneous which opens the possibility for selecting a sub-region with nondisplaceable tracer uptake as a reference region. Future studies in larger animals will likely identify additional regions suitable for utility as a reference region for kinetic modeling. Otherwise, arterial blood sampling will be needed to obtain an input function to support compartmental modeling methods.

### Radiometabolism of [^18^F]-NT160

The extent of radiometabolism of [^18^F]-NT160 in the rat brain and plasma at 60 minutes post injection is shown in the high-performance liquid chromatography (HPLC) chromatogram (Figure 7). Analytical HPLC revealed that the parent [^18^F]-NT160 remained mostly intact (83.0 ±3%) in the brain during the entire scan time of 60 minutes post injection (Figure 7A). In addition to the parent [^18^F]-NT160, two polar radiometabolites M1 (6.0 ±1%) and M2 (11.0 ±3%) were also present in brain homogenates albeit in relatively smaller quantities (Figure 7A). HPLC analysis of the plasma at 60 minutes post injection showed a smaller fraction of the parent [^18^F]-NT160 (30 ±2%) and a single polar radiometabolites (70 ±2%) in which the elution time matches the M1 radiometabolite observed in the brain (Figure 7B). The lack of M2 in the plasma suggests that M2 was produced by radiometabolism in the brain. This also indicates that M1 likely originated in the blood and crossed the BBB and accumulated in the brain. Therefore, it was imperative to investigate and decipher the identity of M1 to ensure that the presence of M1 does not confound the PET data quantification by contributing to the class-IIa HDAC signal.

**Figure 7.**
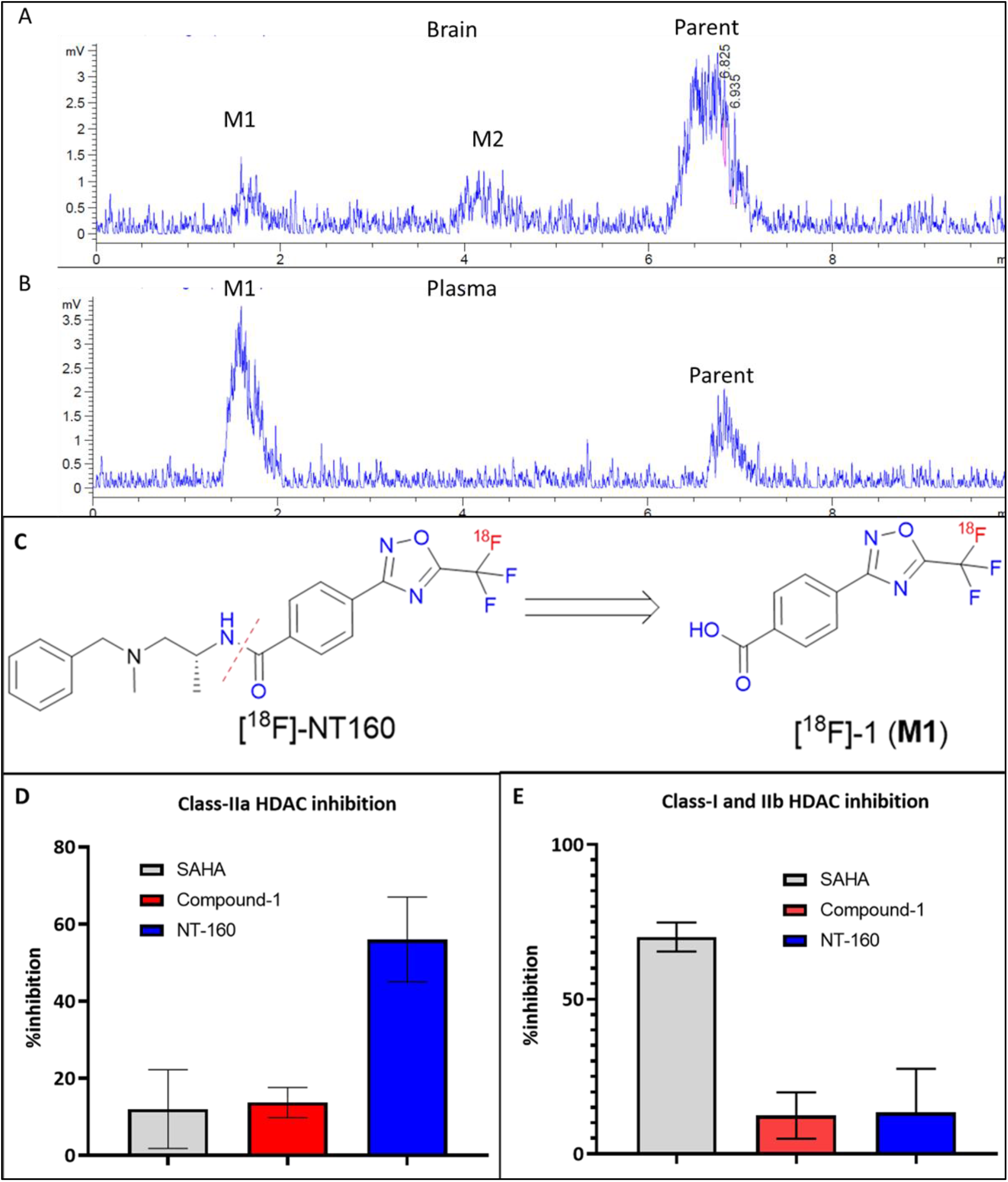
Analytical HPLC chromatograms of [^18^F]-NT160 obtained from A) brain homogenates and B) plasma. [^18^F]-NT160 was eluted with 70% acetonitrile/ammonium acetate buffer (20.0 mM) at flow rate of 1.0 mL/min. C) Proposed blood metabolic pathway for M1 (radiometabolism of [^18^F]-NT160 leads to [^18^F]-1 (M1). D) and E) Inhibition of HDAC activity by compound 1 (M1) in HT-29 cells: D) inhibition of class-IIa HDAC and E) inhibition of class-I/IIb. Compound 1 was screened side-by-side with SAHA and NT160.

**Figure 8.**
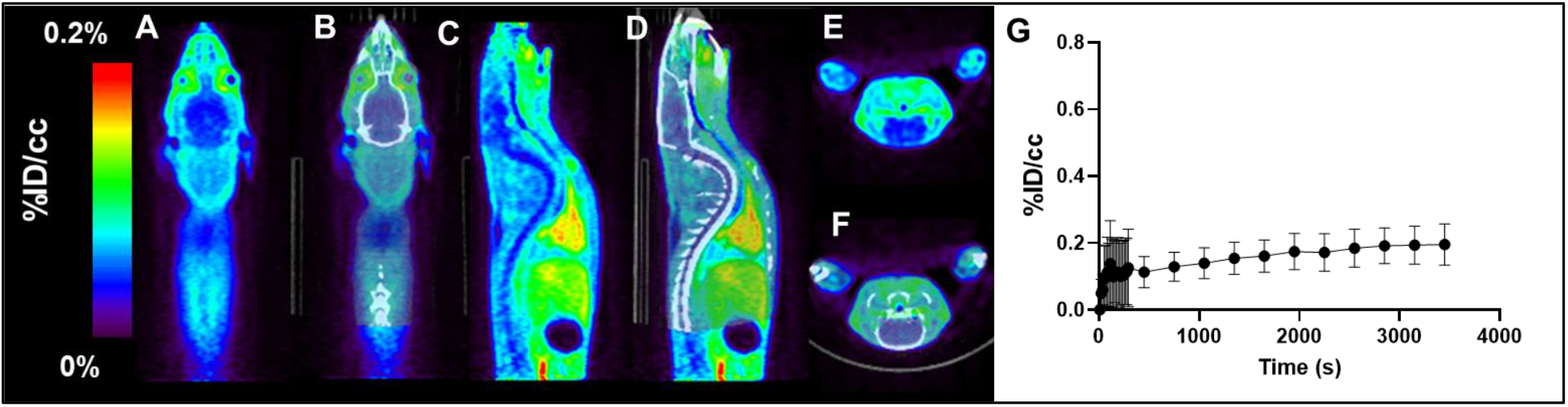
Representative PET, CT, and PT/CT image (summed, 0-60 min) with [^18^F]-1, A) and B) coronal; C) and D) sagittal and E) and F) transverse respectively. G) Time–activity curve of [^18^F]-1 in rat brain.

We surmised that M1 arises from the cleavage of the amide bond thus producing 4-(5-[^18^F]-(trifluoromethyl)-1,2,4-oxadiazol-3-yl)benzoic acid ([^18^F]-**1**) (Figure 7A). We then investigated whether the nonradioactive 4-(5-(trifluoromethyl)-1,2,4-oxadiazol-3-yl)benzoic acid (**1**) can inhibit HDACs using the cell-based assay similar to our previous report ^28^. Our data demonstrate that the IC50 of compound **1** is >10.0 uM (Figure 6 B-C and Figure S.I) for all HDAC classes and thus compound **1** is not an effective inhibitor of HDACs. Compound **1** exhibited poor inhibition (~10%) of class-I, class-IIb and IIa HDACs activities (Figure 7B-C) when compared side-by-side with SAHA (panHDAC inhibitor with higher selectivity to class-I and class-IIb HDACs ^39^) and the parent NT160 (selective class-IIa HDAC) ^28^. The weak affinity of compound **1** to class-I and class-IIb HDACs is highly desirable, since off target binding to these proteins that are expressed in the brain ^40, 41^ may complicate the PET data quantification.

Next, we produced [^18^F]-1 in two radiosynthetic steps similar to the radiosynthesis of [^18^F]-NT160 and our previous report ^29^ (details are provided in the S.I.). Dynamic PET imaging obtained with [^18^F]-1 for 60 minutes revealed that [^18^F]-1 can cross the BBB albeit in relatively low brain uptake of 0.2 %ID/cc (SUV= 0.4) (Figure 7) compared to [^18^F]-NT160. Critically, [^18^F]-1 displayed slow pharmacokinetics and uniform uptake in the brain and therefore is not expected to contribute to the regional biodistribution of [^18^F]-NT160, however, [^18^F]-1 (M1) is expected to slightly increase the overall volume of distribution (V_T_) in the whole brain. The Time–activity curve of [^18^F]-1 (M1) further confirmed the relatively low brain uptake of [^18^F]-1 compared to the parent [^18^F]-NT160. HPLC and R-TLC analysis showed that the parent [^18^F]-1 alone was present in the brain homogenates at 30 minutes post injection. Further analysis by radioactive thin layer chromatography (R-TLC) indicates that the [^18^F]-1 is in fact the polar metabolite (M1) that was observed in brain homogenates of [^18^F]-NT160. (Figure 5: S.I.).

It is important to note that the absolute brain uptake of [^18^F]-1 released from the parent [^18^F]-NT160 is likely to be much less than that of the free [^18^F]-1. [^18^F]-NT160 exhibited fast clearance from blood which is expected to substantially reduce the amount of [^18^F]-1 in circulation and thus reduces the contribution of [^18^F]-1 to the PET signal in the brain compared to the free [^18^F]-1. Deciphering the structures of M1 and M2 will be critical to design the next generation of class-IIa HDAC PET tracers by selecting additional radiolabeling sites (i.e., repositioning of the [^18^F]fluoride or by incorporating [^11^C] radionuclide).

The above data indicates that [^18^F]-1 (M1) is not expected to contribute to the specific binding of class-IIa HDAC or the non-specific and off target binding to other HDACs *in vivo.* [^18^F]-1 (M1) displayed uniform uptake across the brain and therefore, the presence of [^18^F]-1 (M1) in the brain is of less concern as a reference tissue model can be reliably applied ^42^. Moreover, [^18^F]-1 displayed a uniform distribution in the brain, despite containing both the TFMO and the linker pharmacophores of [^18^F]-NT160, thus further demonstrating and supporting the specificity of [^18^F]-NT160 to class-IIa HDACs in the brain. The identity of M2 has not been determined yet, however, it is likely to have arisen in part from metabolism by de-methylation of the methyl amine of [^18^F]-NT160. Further studies are needed to confirm this speculation. However, the absence of M2 in the plasma and the presence of M2 solely in the brain in relatively smaller quantity, indicates that M2 is unlikely to confound the PET data quantification.

## Conclusions

In summary, we reported a **4**-hydroxy-TEMPO assisted radiosynthesis of [^18^F]-NT160, a radioligand for molecular imaging of class-IIa HDACs, a key component of the epigenetic machinery. PET imaging studies with [^18^F]-NT160, a highly potent and selective class-IIa HDACs inhibitor demonstrated a favorable profile for *in vivo* imaging and is useful CNS lead PET tracer. [^18^F]-NT160 displayed excellent pharmacokinetic and imaging characteristics: brain uptake is high in gray matter regions, leading to high-quality PET images; tissue kinetics are appropriate for an [^18^F]-tracer and specific binding for class-IIa HDAC is demonstrated by self-blockade *in vivo* and *in vitro* using autoradiography. *In vivo* molecular imaging with [^18^F]-NT160 provided the first map of distribution of class-IIa HDAC density in the brain and therefore our studies will likely pave the way for accurate comparison of class-IIa expression in disease and health state. However, the moderate specific activity of the [^18^F]-NT160 is a limitation due to the presence of a high mass dose of NT160 that may affect occupancy of class-IIa HDAC in the brain and the periphery and thus can impact the overall brain uptake and may elicit pharmacological response. Therefore, further modifications of [^18^F]-NT160 by repositioning the radiolabel to overcome the inherent moderate specific activity are ongoing and will be reported in due course.

### Methods

#### General information

Solvents and starting material were obtained from commercial sources and were used as received. High-Performance Liquid Chromatography (HPLC) was performed with a 1260 series pump (Agilent Technologies, Stuttgart, Germany) with a built-in UV detector operated at 254 nm and a radioactivity detector with a single-channel analyzer (labLogic) using a semipreparative C18 reverse-phase column (10×250 mm, Phenomenex) and an analytical C18 column (4.6×250 mm, ASCENTIS RP-AMIDE, Sigma). An acetonitrile/ammonium acetate buffer (MeCN/NH_4_OAc: 20 mM) solvents was used for quality control analyses at a flow of 1 mL/min.

#### Chemistry and Radiochemistry

The synthesis NT160 was performed according to our previous report^29^.

#### Radiosynthesis of [^18^F]-NT160

The [18F] was trapped on a QMA cartridge and then eluted to a V-vial (Wheaton) with 80% acetonitrile and 20% water (1.0 mL) solution that contains kryptofix2.2.2 (12.0 mg) and Cs_2_CO_3_ (1.0 mg). The solvent was removed under a stream of Argon at 110 °C. Water residue was removed azeotropically with the addition of acetonitrile (3 x 1.0 mL) and repeated drying under a stream of Argon at 110 °C.

The bromo-precursor (6-8 mg) mixed with 4-hydroxy-TEMPO (0.3-0.5 mg) was dissolved in DMSO (0.4 mL) by heating at 110 °C for 2 minutes. The solution was added to the dried Cs[18F]/kryptofix2.2.2 and the mixture was heated at 165 °C for 20 minutes. The reaction mixture was cooled at room temperature in a water bath for 2 minutes and was then diluted with 30% methanol in dichloromethane (2.5 mL) passed through a silica gel cartridge (waters, 900 mg). The silica cartridge was further washed with 1.0 mL of methanol and the solvent were evaporated under a stream of argon at 60-80 °C. The residue was redissolved in a solution (1.5 mL) containing 30% acetonitrile and 70% ammonium acetate (NH_4_OAc: 20 mM). Purification was performed using semipreparative high-performance liquid chromatography (HPLC) with a 1260 series pump (Agilent Technologies, Stuttgart, Germany) with a built-in UV detector operated at 250 nm and a radioactivity detector with a single-channel analyzer (labLogic). [^18^F]-NT160 was eluted from C18 reverse-phase column (10×250 mm, Phenomenex) using 67% acetonitrile: 33% ammonium acetate buffer (NH_4_OAc: 20 mM) at flow rate of 4.0 mL/minute. [^18^F]-NT160 was eluted and collected at 19-21 min post injection. Water (15.0 mL) was added to the solution and the solution was trapped on a C-18 light cartridge and eluted with ethanol (0.3 mL) and formulated for *in vivo* and in *vitro* studies.

#### Radiosynthesis of [^18^F]-1

The details of the radiosynthesis of [^18^F]-1 is provided in the supplementary material.

#### HDAC Inhibitor Biochemical Assay

The details of the cell-based assay are provided in the supplementary material. The cell-based HDAC inhibitor assays were performed in HT-29 cells (ATCC). Briefly, cells were grown in DMEM media (Gibco) supplemented with 10% FBS (Thermo Fisher Scientific) and 1X Anti-Anti (Gibco) in a humidified incubator at 37°C with 5% CO_2_. The day before the experiment, cells were washed with PBS and dissociated using 0.25% Trypsin-EDTA (Gibco). Trypsinization was stopped with media containing serum, and cells were collected by centrifugation. The cells were then washed with HDAC Assay Buffer (RPMI 1640 media without phenol red (Gibco) containing 0.1% FBS), re-collected by centrifugation, resuspended in HDAC Assay Buffer, and counted. Two hundred thousand cells were seeded into 96 well plates and placed into the incubator overnight. HT-29 cells (obtained from ATCC) with the addition of either 100 μM Boc-Lys-TFA (class-IIa selective substrate) or 200 μM Boc-Lys-Ac (class-I/IIb selective substrate) were plated into 96 well plates at 200,000 cells/well in 45 uL cellular assay buffer (RPMI without phenol red, 0.1% Fetal Bovine Serum) and were incubated for 3 h at 37°C. The deacetylation achieved by the addition of 50 uL HDAC developer solution (2.5 mg/ml trypsin in DMEM without Fetal Bovine Serum and 10% tween 80), followed by incubation for one hour to sensitize the substrate and to lyse the cells. Fluorescent counts were read with microplate reader at an excitation wavelength of 360 nm and detection of emitted light of 460 nm.

#### PET Imaging Procedures in Animals

All studies were performed under a protocol approved by the Institutional Animal Care and Use Committee of Stony Brook University. *In vivo* microPET/CT imaging studies were performed in healthy rats as described below. Anesthetized Sprague Dawley rats (200-450 g, N=6) were placed in the Inveon uPET (Siemens, Knoxville, TN) in the supine position with the skull positioned in the center of the field of view. [^18^F]-NT160 (18.5-25.9 MBq/animal) was administered via the tail-vein injection in a total volume 0.5-1. 2 ml. Dynamic PET images were obtained over 60 minutes followed by CT imaging (15 min scan). Images were reconstructed with attenuation correction using an ordered subset expectation maximization (OSEM2D) algorithm with 16 subsets and 4 iterations. PET image analysis was performed using the AMIDE software by using manual segmentation of regions of interest (ROI), including whole brain, cortex, hippocampus, thalamus, cerebellum, and brain stem. Rat Brain Atlas was used for alignment and identification of specific anatomical markers of the brain^35, 36^. The time-activity curve (TAC) was generated from the region of interest (ROI) by plotting the radioactivity/cc vs time. Levels of accumulation of the radiotracer in tissues were expressed as standard uptake values (SUV) that were calculated for the regions of interest (ROI) using the AMIDE software. The SUV is defined as the ratio of the tissue radioactivity concentration C (e.g., expressed as Bq/g tissue) at given time point post injection T, and the injected dose (e.g., in Bq, decay-corrected to the same time T), and normalized by the body weight in grams. GraphPad Prism (Graph Pad Software La Jolla, CA) was used for image data analysis.

#### *In vitro* Autoradiography

Adult rats were sacrificed, and the brains were fixed in 4%PFA paraffin overnight, then embedded into paraffin block. Brains were sectioned into slices (10 um). Then the slices were deparaffinized in xylene (three times) for 30 minutes (three times) followed by rehydration with 100% ethanol 95% ethanol 70 % ethanol and water for 30 minutes followed by rinsing in R.O water for at least 30 minutes. The brain slides used for blocking studies were incubated with NT-160 in 20 ml of 1 uM solution of NT-160 for 1 hour then preincubated and washed with binding buffer (50 mM Tris base, 2 mM MgCl2, pH 7.4) with 0.1% BSA for 10 min. ^18^F-NT160 (110.0 μCi) was added in 20 ml binding buffer and incubated at room temperature for 120 min. Subsequently, the slides were washed twice for 10 minutes in ice-cold binding buffer and dipped in ice-cold water before being dried. The slides were placed onto the storage phosphor screen in the dark overnight

#### Histology

Rat brains were fixed in 4%PFA paraffin overnight, then embedded into paraffin block. Brains were sectioned into slices (10 um). The slices were deparaffinize with xylene for 30 minutes three times, rehydrated with 10%ethanol, 90% ethanol, 70% ethanol and water consecutively. Antigen retrieval was performed with citrate buffer (pH=6) overnight at 60C° followed by washing with PBS (three times). The slices were permeabilized with 0.1% Triton X-100 in 2.5% goat serum and stained with primer HDAC4 antibody (ABCAM, ab123513) overnight at 4°C followed by washing with PBS (three times). The slices were incubated with biotinylated secondary antibody for 1 hour, then were washed with PBS (three times). The slices were incubated with ABC agent for 30 minutes. The color was developed with DAB for two minutes followed by washing with water (three times). The staining was counted with hematoxylin for 1 minute.

#### Radiometabolite studies with [^18^F]-NT160

The rat brains from the above imaging studies were excised, hemisected, and half brain was homogenized in 1.0 mL acetonitrile. The resulting suspension was centrifuged, and the clear supernatant was vortexed and injected into analytical radio-HPLC and the extent of metabolism in the brain was determined. The radioactive peaks were compared to the retention time of the parent [^18^F]-NT160. Also, samples were collected at a fraction of 1.0 minutes and counted using a Gamma Counter. The %radioactivity of the new peaks compared to the %radioactivity of the peak of the parent tracer was used to determine the extent of radiolabeled metabolites in the brain.

Blood was collected, and plasma was obtained by centrifugation. Protein-free plasma was obtained by addition of acetonitrile followed by centrifugation. The acetonitrile fraction was injected into analytical radio-HPLC (67% acetonitrile: ammonium acetate buffer (20mM) solution). The radioactive peaks were compared to the retention time of the parent [^18^F]-NT160 (~6.9 minutes) ^28^. Also, samples were collected at a fraction of 1.0 minutes and counted using a Gamma Counter. The %radioactivity of the new peaks compared to the %radioactivity of the peak of the parent tracer was used to determine the extent of radiolabeled metabolites in the blood.

