## Supplementary material for "High-Contrast PET imaging with [^18^F]-NT160, a Class-IIa Histone Deacetylase (Class-IIa HDAC) Probe for In Vivo Imaging of Epigenetic Machinery in the Central Nervous System": SI

**Quality control analyses:** The identity of [$^{18}$F]-NT160 was confirmed using analytical C18 column (4.6×250 mm, ASCENTIS RP-AMIDE, Sigma) by co-eluting with authentic solution of NT160 using 70% acetonitrile: 30% ammonium acetate buffer (NH4OAc: 20 mM) at flow rate of 1.0 mL/minute. Radiochemical purity was determined by integrating the radioactive peak corresponding to [$^{18}$F]-NT160 against all other radioactive peaks present in the HPLC chromatograms. [$^{18}$F]-NT160 was obtained reproducibly in high radiochemical purity (>98%). [$^{18}$F]-NT160 stability in the formulated solution was examined for four hours by measuring and comparing the integration additional (none-observed) radioactive peaks at 1, 2, 3 and 4 hours post formulation to the radioactive peak corresponding to [$^{18}$F]-NT160 obtained at the end of the experiment. No additional radioactive peaks were observed thus verifying the radiochemical stability of [$^{18}$F]-NT160.

**Molar activity:** The molar activity was determined from the area under the curve of the tracer that is attributed to the ultraviolet peak in the HPLC chromatogram (Figure 1) against a calibration curve pre-prepared with the unlabeled reference standard.

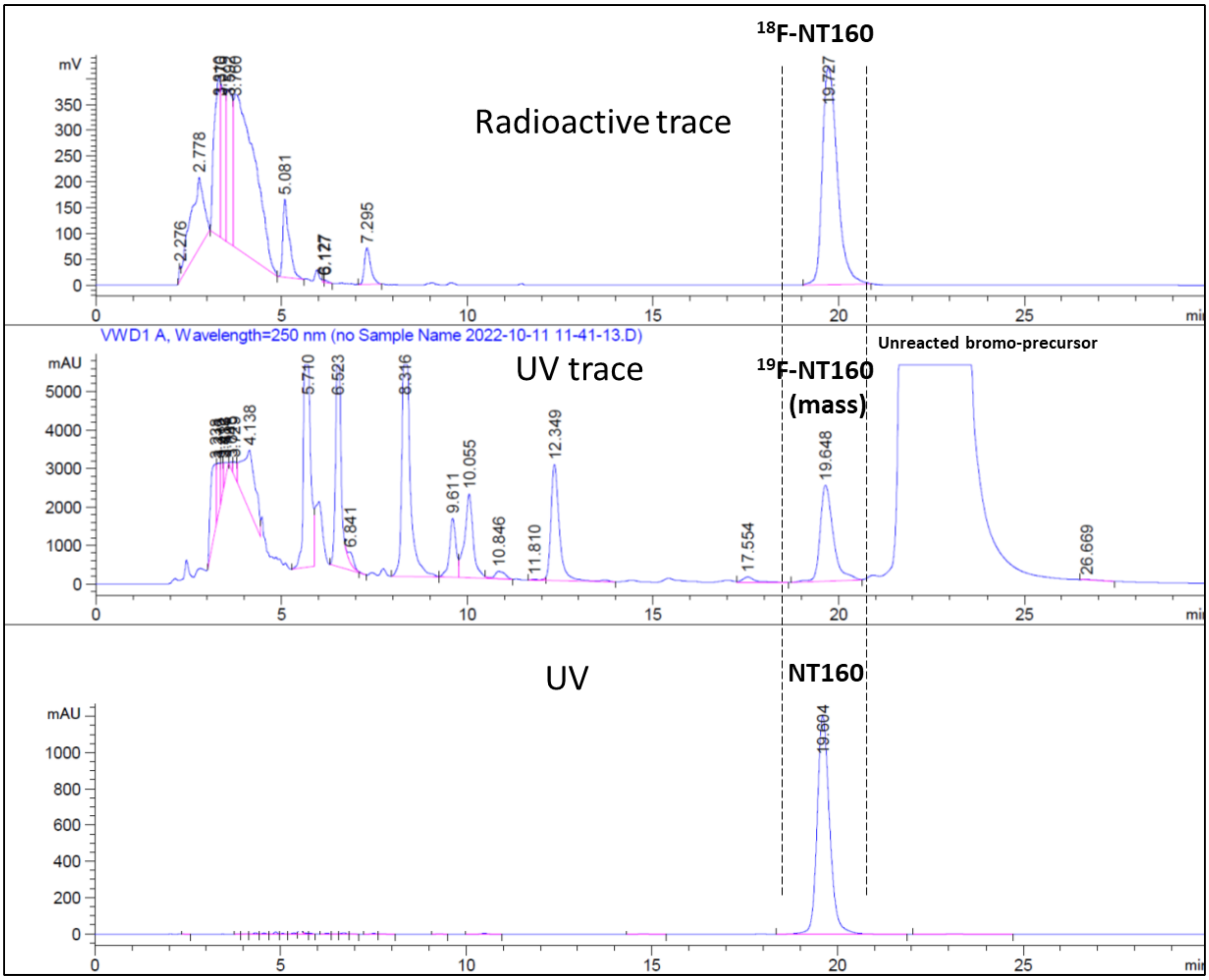


**Figure 1.** Semi-preparative HPLC chromatograms of A) radioactive tracer for purification of [$^{18}$F]-NT160, B) ultraviolet tracer for purification of [$^{18}$F]-NT160 (note the mass related to NT160) and C) ultraviolet chromatogram obtained from authentic sample of NT160. Samples were eluted with 67% acetonitrile/ammonium acetate buffer (20.0 mM)) at a flow rate of 4.0 mL/min.

### Radiosynthesis of [18F]-1

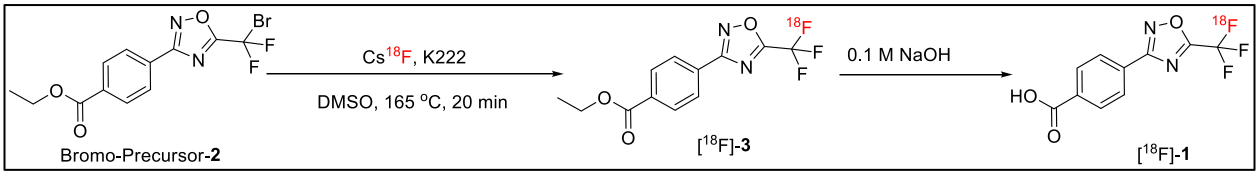


**Scheme 1.** Radiosynthesis of [$^{18}$F]-**1.**

The synthesis of the bromo-precursor-2 was performed according to our previous report [1].

The radiosynthesis of [$^{18}$F]-**3** was performed similar to the radiosynthesis of [$^{18}$F]-NT160 described in the main text. **[$^{18}$F]-3** was purified using 70% acetonitrile: 30% ammonium acetate buffer (NH4OAc: 20 mM) at a flow rate of 4.0 mL/minute. [$^{18}$F]-**3** was collected and trapped on a C-18 light cartridge (Waters) and eluted with ethanol (0.3 mL) and treated with 0.1 N NaOH (0.1 mL) for 10 minutes in which hydrolysis was complete (HPLC analysis). [$^{18}$F]-**1** solution was neutralized with HCl (0.1 mL) and formulated by addition of sterilized water to a <10% concentration of ethanol.

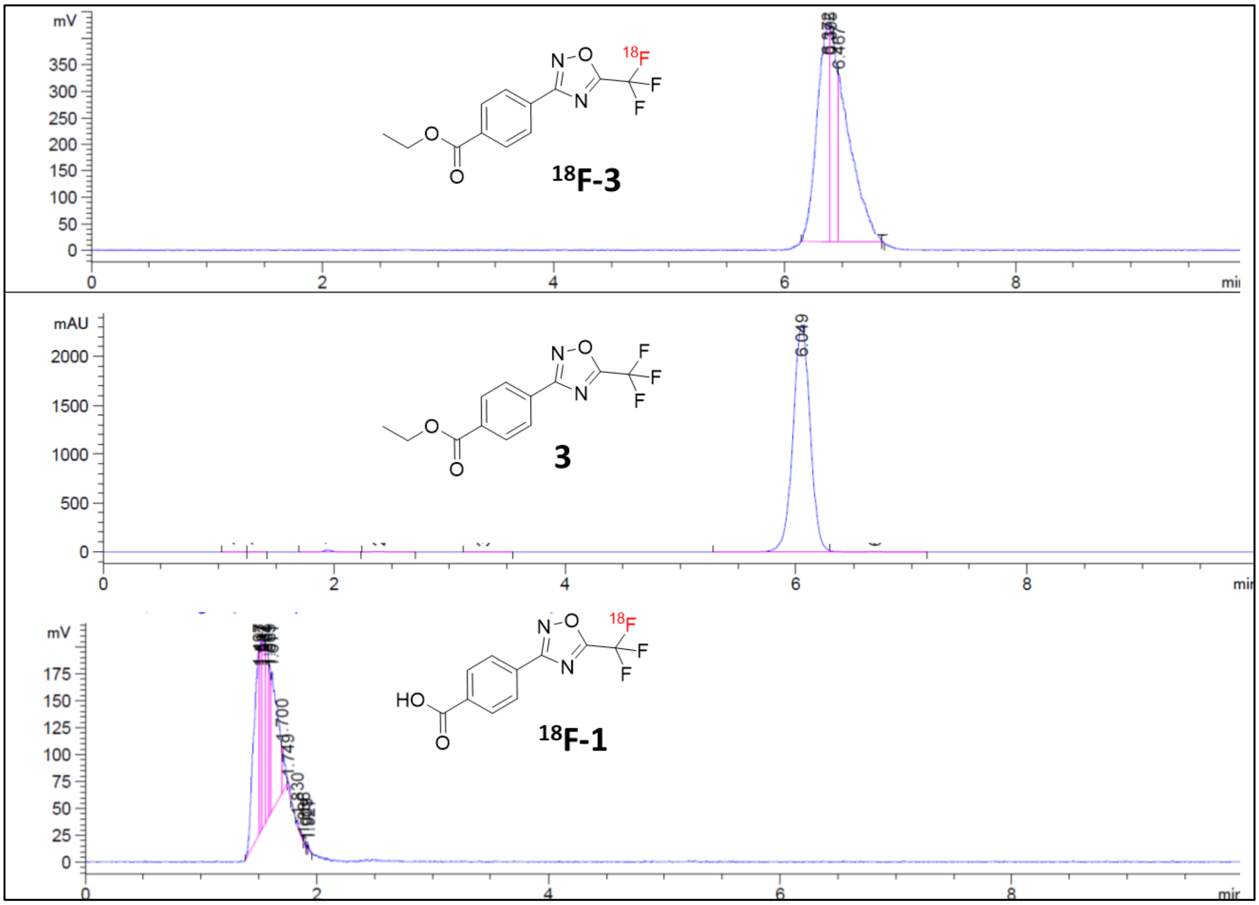


**Figure 2.** Analytical HPLC chromatograms of A) radioactive tracer for purification of [$^{18}$F]-3, B) ultraviolet chromatogram obtained from authentic sample of 3, C) [$^{18}$F]-1 obtained from hydrolysis of [$^{18}$F]-3. Samples were eluted with 70% acetonitrile/30% ammonium acetate buffer (20.0 mM)) at a flow rate of 1.0 mL/min.

### HDAC Inhibitor Biochemical Assay

Cell-based HDAC inhibitor assays were performed in HT-29 cells (ATCC). Briefly, cells were grown in DMEM media (Gibco) supplemented with 10% FBS (Thermo Fisher Scientific) and 1X Anti-Anti (Gibco) in a humidified incubator at 37°C with 5% $CO_2$. The day before the experiment, cells were washed with PBS and dissociated using 0.25% Trypsin-EDTA (Gibco). Trypsinization was stopped with media containing serum, and cells were collected by centrifugation. The cells were then washed with HDAC Assay Buffer (RPMI 1640 media without phenol red (Gibco) containing 0.1% FBS), re-collected by centrifugation, resuspended in HDAC Assay Buffer, and counted. Two hundred thousand cells were seeded into 96 well plates and placed into the incubator overnight.

Stock solutions were prepared ahead of time. HDAC inhibitor stock solutions at 20 mM were made in DMSO (Sigma) and stored in single-use aliquots at -20°C. The HDAC inhibitors include: suberoylanilide hydroxamic acid (SAHA), 5-trifluoromethyl-1,2,4-oxadiazole (TFMO) and NT-160. For the positive inhibitor control, stock solutions of 5 mM Trichostatin A (Tocris) were made in DMSO and stored in single-use aliquots at -20°C. 1 mg/ml aliquots of BSA (Sigma) were made in HDAC Assay Buffer and stored in single-use aliquots at -20°C. Finally, 100 mM stock solutions of the substrate Boc-Lys (Tfa)-AMC and Boc-Lys (Ac)-AMC (Bachem) in DMSO were stored at -20°C and thawed as needed.

The next day, inhibitors were thawed and serially diluted with HDAC Assay Buffer into microcentrifuge tubes. From these dilutions, working solutions were prepared containing a final concentration of 50, 10, or 1 µM inhibitor. For Trichostatin A, the final working concentration was 10 µM. The final working concentration for the class IIa HDAC substrate, Boc-Lys (Tfa)-AMC, was 100 µM; the class I and IIb substrate, Boc-Lys (Ac)-AMC, final concentration was 200 µM.

The working solutions containing inhibitors, controls and substrates were prepared as follows. The final working solution volume per well was 50 µl. Note that 3 technical replicates were performed for each condition. Therefore, working solution containing 200 µl was made for each condition (leaving some extra). The final concentration of DMSO in all samples was maintained at 0.45%. If required, working solutions were supplemented with HDAC Assay Buffer containing 2% DMSO. First, the appropriate volume of HDAC Assay Buffer and 2% DMSO was added to tubes followed by a stock solution of BSA (1 mg/ml) for a final concentration of 0.1 mg/ml. The tubes were closed and mixed thoroughly to coat the tube walls with BSA (to prevent substrate sticking to the tube walls). A microcentrifuge was used to collect the solution at bottom of tubes. Next, the appropriate inhibitor was added to each tube followed by the substrate. The tubes were mixed well and spun again to collect solution at bottom of the tubes. The negative control (no inhibitor) contained substrate and DMSO only at a final concentration of 0.45% in HDAC Assay Buffer. The no HDAC enzyme control (no cells) also contained substrate and 0.45% DMSO; note that this sample is used to subtract background noise from the sample.

Once the working solutions were ready, a P1000 pipette tip was used to carefully remove the media from the appropriate wells of the 96 well plate containing HT-29 cells. Working one sample at a time, the media was removed from the wells and replaced with 50 µl of the working solution. When done, the plate was carefully swirled to mix. This was done with the lid on, and with two hands. The plate was returned to the incubator for 3 hours.

During the incubation period, Developer Solution (2.5 mg/ml Trypsin in RPMI 1640 without phenol red, 10% Tween 20) was made and kept at room temperature. A shaking platform was warmed to 37°C prior to the end of the incubation period. 50 µl of Developer solution was added to each well while making sure there were no bubbles were present which can interfere with subsequent measurements. With the lid on, the plate was placed in a pre-warmed shaker set at 80 rpm for 10 minutes. After 10 minutes, the plate was parafilmed, covered in aluminum foil and stored at 4°C overnight. The next day, fluorescence was measured in a Synergy H1 plate reader. Each sample was read in triplicate. The first two reads were read with the following parameters: 340 nm excitation; 435 nm emissions; Gain ,100; Delay, 100 msec; Read Height, 7 mm, Read Speed, normal; 10 Measurements/Data Point. All parameters were identical for the third read except for the emissions, which were measured at 440 nm. The 3 individual reads/sample were averaged for a final relative fluorescent unit (RFU)/sample.

Four independent experiments (n=4) were performed for this assay. The mean RFU value for each sample was calculated and converted to percent inhibition (relative to DMSO) for each independent experiment. The mean percent inhibition was calculated for all 4 independent experiments. Standard Deviation (SD) and the Standard Error of the Mean (SEM) were calculated for each sample. Error bars on the graphs for each substrate represent SEM.

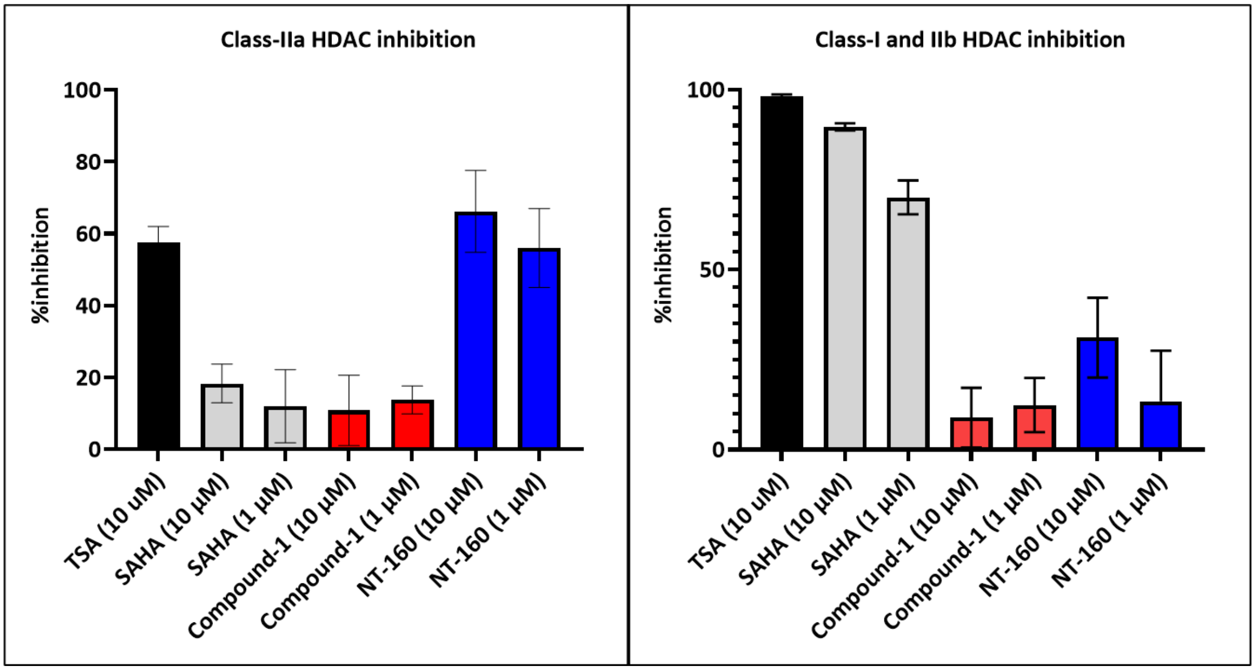


**Figure 3.** Cellular (HT-29 cells) HDAC inhibition by compound 1 (M1) of class-IIa HDAC and class-I/IIb respectively, screened side-by-side with SAHA, TSA and NT160 at 1.0 and 10.0 uMol.

### *In Vivo* Blocking Studies:

The *in vivo* displacement studies were performed similar to the PET imaging studies except that the animals were injected (i.v.) with 500 ug/kg of the NT160, 10 min prior to the *in vivo* imaging. SUV was used to quantify the extent of pretreatment with NT160 on the brain.

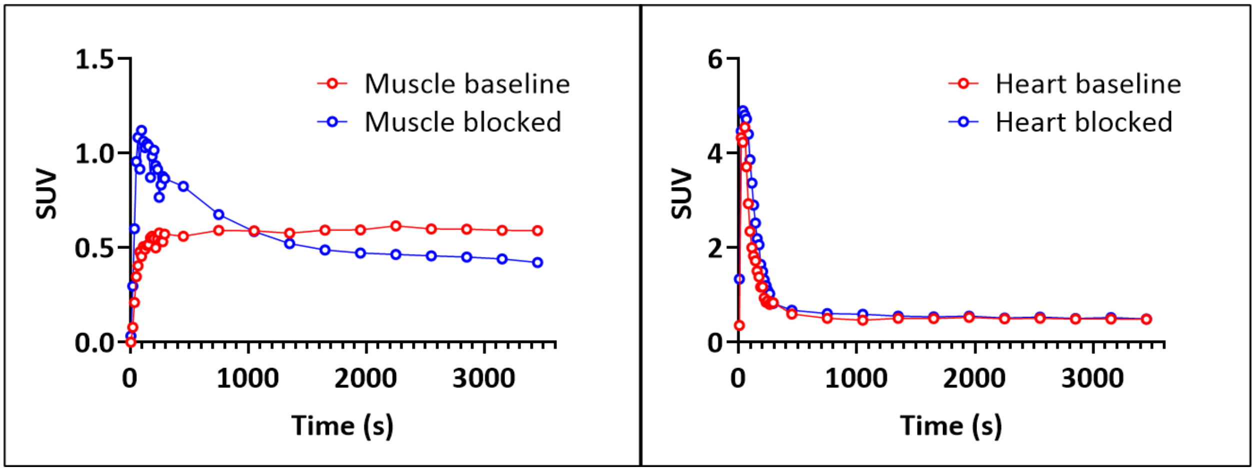


**Figure 4.** Time–activity curves of [$^{18}$F]-NT160 obtained from dynamic imaging for 60 min. A) muscle, and heart at baseline and B0 after pretreatment with NT160 (0.5 mg/kg).

***Ex vivo* imaging at baseline and after self-blocking**. Rats (N=2/study group) were injected with 200 uCi of [$^{18}$F]-NT160 alone (baseline) or with a dose of the non-radioactive NT160 (1.0 mg/kg). At 20 min post injection, animals were killed, and their brains were excised and were imaged side-by-side ex-vivo. The brain uptake of $^{18}$F-NT160, was obtained by counting the radioactivity (CPM: count per minute) using gamma counter and normalized to the brain weight. Static PET imaging showed that self-blocking with the unlabeled NT160 resulted in decrease in the total tracer uptake in the brain: 30% decrease (Figure 5A-D) thus demonstrating specific binding of [$^{18}$F]-NT160 to class-IIa HDACs. Also, the brain uptake was reduced when comparing radioactivity/weight (CPM/g) obtained from the brain with tracer alone (baseline) and with the tracer co-administered with NT160. Therefore, the blocking study confirmed the specific binding of the tracer to class-IIa HDACs in the brain.

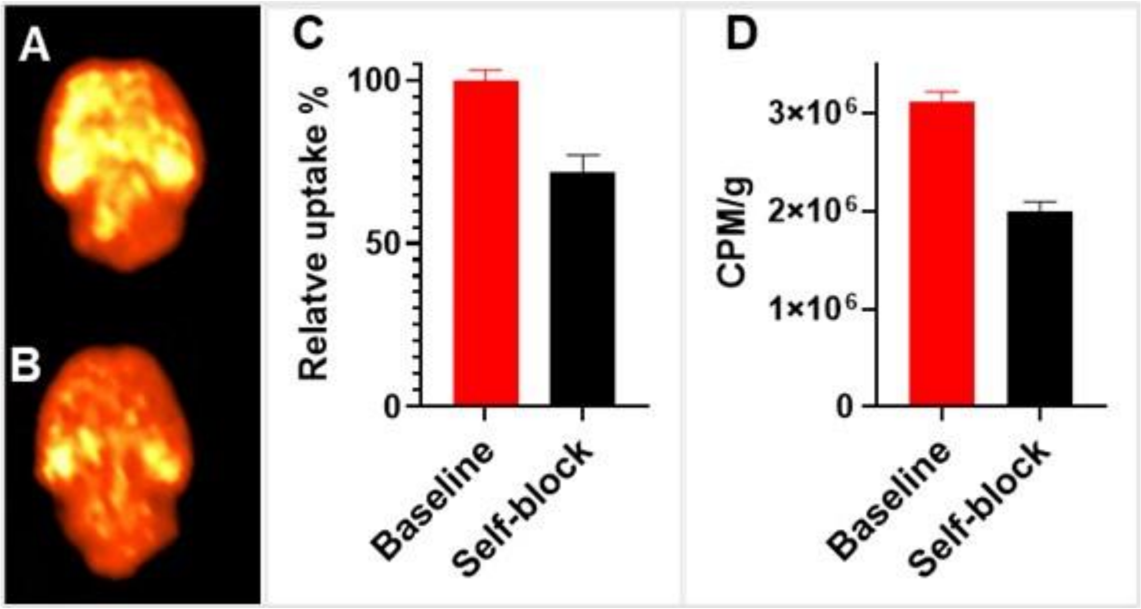


**Figure 5.** Side-by-side *ex-vivo* PET imaging of rat brain at 20 min after injection of $^{18}$F-NT160 A) baseline and B) after treatment with NT160 (1.0 mg/kg). C) Self-blocking shows reduced SUV by 30% of [$^{18}$F]-NT160 in rat brain with the non-radioactive NT160 and D) self-blocking shows reduced brain uptake by 32% using gamma counting.

### Radiometabolite studies with [18F]-NT160:

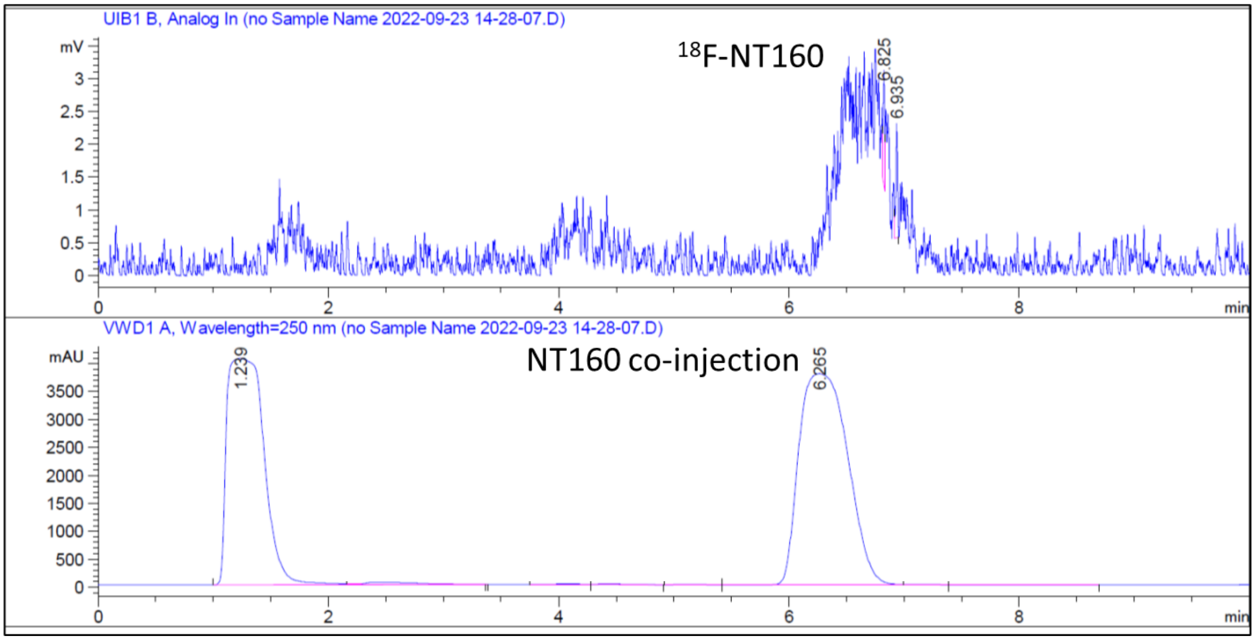


**Figure 5.** HPLC chromatograms of [$^{18}$F]-NT160 brain radiometabolites co-injected with authentic NT160. Samples were eluted with 70% acetonitrile/ammonium acetate buffer (20.0 mM)) at a flow rate of 1.0 mL/min.

### Radiometabolite studies with [18F]-1(M1):

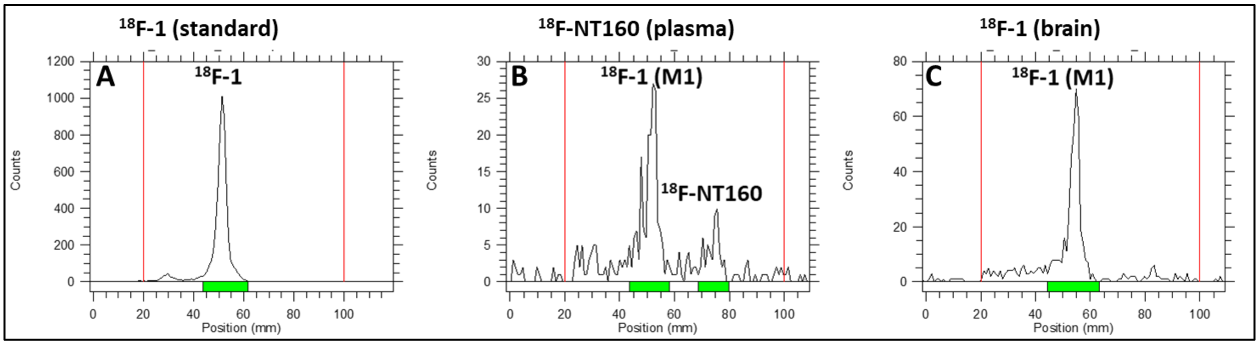


**Figure 6.** Radioactive thin layer chromatography (R-TLC) analysis: A) [$^{18}$F]-1 alone, B) brain radiometabolites of [$^{18}$F]-NT160 and C) [$^{18}$F]-1 in brain homogenates. Note the $R_f$ value of [$^{18}$F]-1 is consistent among the three experiments thus confirming the M1 is at least in part is [$^{18}$F]-1. Condition: methanol (8%): ethyl acetate (92%).

1. Turkman, N.; Liu, D.; Pirola, I., Novel late-stage radiosynthesis of 5-[18F]-trifluoromethyl-1,2,4-oxadiazole (TFMO) containing molecules for PET imaging. *Sci Rep* **2021,** *11* (1), 10668.
